# The mitochondrial surface receptor Tom70 protects the cytosol against mitoprotein-induced stress

**DOI:** 10.1101/2020.09.14.296194

**Authors:** Sandra Backes, Yury S. Bykov, Markus Räschle, Jialin Zhou, Svenja Lenhard, Lena Krämer, Timo Mühlhaus, Chen Bibi, Cosimo Jann, Justin D. Smith, Lars M. Steinmetz, Doron Rapaport, Zuzana Storchová, Maya Schuldiner, Felix Boos, Johannes M. Herrmann

## Abstract

Most mitochondrial proteins are synthesized as precursors in the cytosol and post-translationally transported into mitochondria. The mitochondrial surface protein Tom70 acts at the interface of the cytosol and mitochondria. *In vitro* import experiments identified Tom70 as targeting receptor, particularly for hydrophobic carriers. Using *in vivo* methods and high content screens, we revisited the question of Tom70 function and considerably expanded the set of Tom70-dependent mitochondrial proteins. We demonstrate that the crucial activity of Tom70 is its ability to recruit cytosolic chaperones to the outer membrane. Indeed, tethering an unrelated chaperone-binding domain onto the mitochondrial surface complements most of the defects caused by Tom70 deletion. Tom70-mediated chaperone recruitment reduces the proteotoxicity of mitochondrial precursor proteins, in particular of hydrophobic inner membrane proteins. Thus, our work suggests that the predominant function of Tom70 is to tether cytosolic chaperones to the outer mitochondrial membrane, rather than to serve as a mitochondria-specifying targeting receptor.

## Introduction

With a concentration of 30,000 to 50,000 ribosomes per μm^3^, the eukaryotic cytosol is densely packed with molecular machines for protein synthesis which make up a considerable fraction of its total volume (Marini et al., 2020). Rapid and efficient protein folding in the cytosol is of pivotal importance for rapidly growing cells. Chaperones, particularly those of the Hsp70 and Hsp90 family, with the assistance of different co-chaperones and accessory factors, bind to nascent chains as soon as they emerge from the ribosome in order to facilitate their folding (Hartl et al., 2011; Kramer et al., 2019; Sontag et al., 2017), or hold them in a translocation-competent state for transport across membranes of organelles (Deshaies et al., 1988; Hoseini et al., 2016; Jores et al., 2018; Young et al., 2003).

Mitochondria are essential organelles of eukaryotic cells. They synthesize a handful of very hydrophobic polypeptides on mitochondrial ribosomes in the matrix. All other 900 (yeast) to 1500 (humans) mitochondrial proteins are encoded by nuclear genes and synthesized in the cytosol (Calvo et al., 2016; Morgenstern et al., 2017; Vogtle et al., 2017). With the exception of a small number of inner membrane proteins (Tsuboi et al., 2020; Williams et al., 2014), the import of mitochondrial proteins occurs post-translationally, meaning that they are first synthesized in the cytosol and subsequently translocated into mitochondria; however, the spatial-temporal details of these processes are largely elusive (Gold et al., 2017; Jan et al., 2014).

Mitochondrial protein biogenesis strictly depends on the cytosolic chaperone capacity (Becker et al., 1996; Ben-Menachem et al., 2018; Deshaies et al., 1988; Doring et al., 2017; Hoseini et al., 2016; Pfanner et al., 1987; Stein et al., 2019; Terada et al., 1996). Presumably as a consequence of their strong tendency to sequester chaperones, precursor proteins accumulating in the cytosol induce a sudden growth arrest, trigger the heat shock response to increase components of the chaperone and proteasome system, and activate specific factors on the mitochondrial surface that clean off translocation intermediates (Boos et al., 2019; Boos et al., 2020; Martensson et al., 2019; Shakya et al., 2020; Wang and Chen, 2015; Weidberg and Amon, 2018; Wrobel et al., 2015).

Most mitochondrial proteins are synthesized with an N-terminal presequence that serves as a matrix-targeting signal (MTS) (Vögtle et al., 2009; von Heijne, 1986). Presequences are recognized by Tom20 and Tom22, two receptor proteins that are part of the translocase of the outer membrane (TOM) complex (Araiso et al., 2019; Rimmer et al., 2011; Shiota et al., 2015) before they lead the way into the matrix, across protein-conducting channels of the TOM complex and the presequence translocase (or TIM23 complex) in the inner membrane (Chacinska et al., 2009). Internal MTS-like (iMTS-L) sequences are frequently found in matrix proteins; though not sufficient as import signals, these patterns can strongly improve the import-competence of precursors (Backes et al., 2018).

Many mitochondrial proteins, however, lack presequences and embark on other import routes into mitochondria. This is the case for all proteins of the outer membrane and most components of the intermembrane space (IMS) which employ several distinct pathways (Doan et al., 2020; Drwesh and Rapaport, 2020; Edwards et al., 2020; Finger and Riemer, 2020; Wiedemann and Pfanner, 2017). In addition, many mitochondrial inner membrane proteins, in particular the members of the metabolite carrier family (carriers for short) lack presequences and are imported via a distinct ‘carrier pathway’ (Horten et al., 2020; Rehling et al., 2004). From studies using the ATP/ADP carrier (*i.e*. Pet9 in yeast) as a model protein, it was proposed that the carrier pathway differs from the import route of matrix-destined proteins already on the surface of mitochondria, where carriers bind the ‘carrier receptor’ Tom70 which would insert them into the universal protein-conducting channel of the TOM complex. In the IMS, a soluble chaperone complex consisting of small Tim proteins further transfers the carriers to a specific translocase of the inner membrane, the TIM22 complex, for membrane insertion. Although the mitochondrial steps of the carrier pathways were dissected in detail by powerful *in vitro* assays (Hasson et al., 2010; Pfanner and Neupert, 1987; Ryan et al., 1999), the early, *i.e*. cytosolic steps remain unclear.

This lack of understanding is particularly obvious for the role of Tom70 (and its paralog Tom71), although this outer membrane protein was one of the first import components discovered (Hines et al., 1990; Söllner et al., 1990; Steger et al., 1990). In contrast to all other TOM subunits, Tom70 is presumably no stoichiometric component of the TOM complex, but rather associates with the outer membrane translocase in a dynamic and transient fashion.

Tom70 offers dedicated binding sites for the recruitment of cytosolic Hsp70 and Hsp90 chaperones via their unique C-terminal EEVD tails (Young et al., 2003) and was found to interact with co-chaperones (Opalinski et al., 2018). It also directly binds mitochondrial precursor proteins (Brix et al., 1999; Brix et al., 2000; Iwata and Nakai, 1998; Melin et al., 2015; Papic et al., 2011). However, the specific contribution of each of these properties in the context of mitochondrial protein import is not clear, particularly, because these functions of Tom70 cannot be reliably assessed with the *in vitro* import assays that were used in most studies.

In this study, we used a number of complementary *in vivo* approaches to elucidate the specific role of Tom70. Our assays demonstrate that Tom70 is not a specific receptor for carriers. Rather, Tom70 supports the biogenesis of some carriers (in particular of Pet9), but also that of many other mitochondrial proteins. Many Tom70 clients contain hydrophobic transmembrane segments or other aggregation-prone regions. Most of these proteins are also sensitive to heat, in line with the temperature-sensitivity of Tom70-deficient cells. Interestingly, the loss of Tom70 can be largely complemented in strains carrying an unrelated EEVD-binding protein on the mitochondrial surface. Thus, the predominant and crucial function of Tom70 is not that of a classical import receptor. Instead, Tom70 serves as co-chaperone on the mitochondrial surface to suppresses the toxic effects of mitochondrial precursor proteins.

## Results

### Tom70 is required for the biogenesis of many mitochondrial proteins

On the basis of *in vitro* experiments with a very small number of substrates, Tom70 was proposed to serve as a mitochondrial import receptor for precursors that are made without presequences (Becker et al., 2011; Brix et al., 1999; Papic et al., 2011; Steger et al., 1990), in particular of carriers such as Pet9 or Oac1 (Fig. 1A). Tom70 also supports the *in vitro* import reaction of some presequence-containing proteins such as Atp1, but not that of others, such as Hsp60 (Fig. 1B). Hence, to date, there is no clear understanding of the substrate range of Tom70. To elucidate the *in vivo* substrate spectrum of Tom70, we hypothesized that proteins that do not get properly imported into mitochondria will be degraded by cytosolic quality control and hence that we can use their steady-state cellular abundance as a proxy for their capacity to be properly targeted to mitochondria. Therefore, we measured, using quantitative mass spectrometry, the levels of all mitochondrial proteins in Δ*tom70/71* mutants lacking the genes for Tom70 and its barely expressed paralog Tom71 (Schlossmann et al., 1996) (Fig. 1C). Cells were grown on galactose at 30°C, at which loss of Tom70 doesn’t result in reduced growth rates, to avoid secondary growth-dependent effects. Cells were harvested at exponential growth phase from which proteins were extracted and digested with trypsin. Peptides were labeled using a tandem mass tag (TMT) protocol and multiplexed to quantify proteins in combined sample runs (samples from additional mutants, described later, were also pooled).

**Fig. 1.**
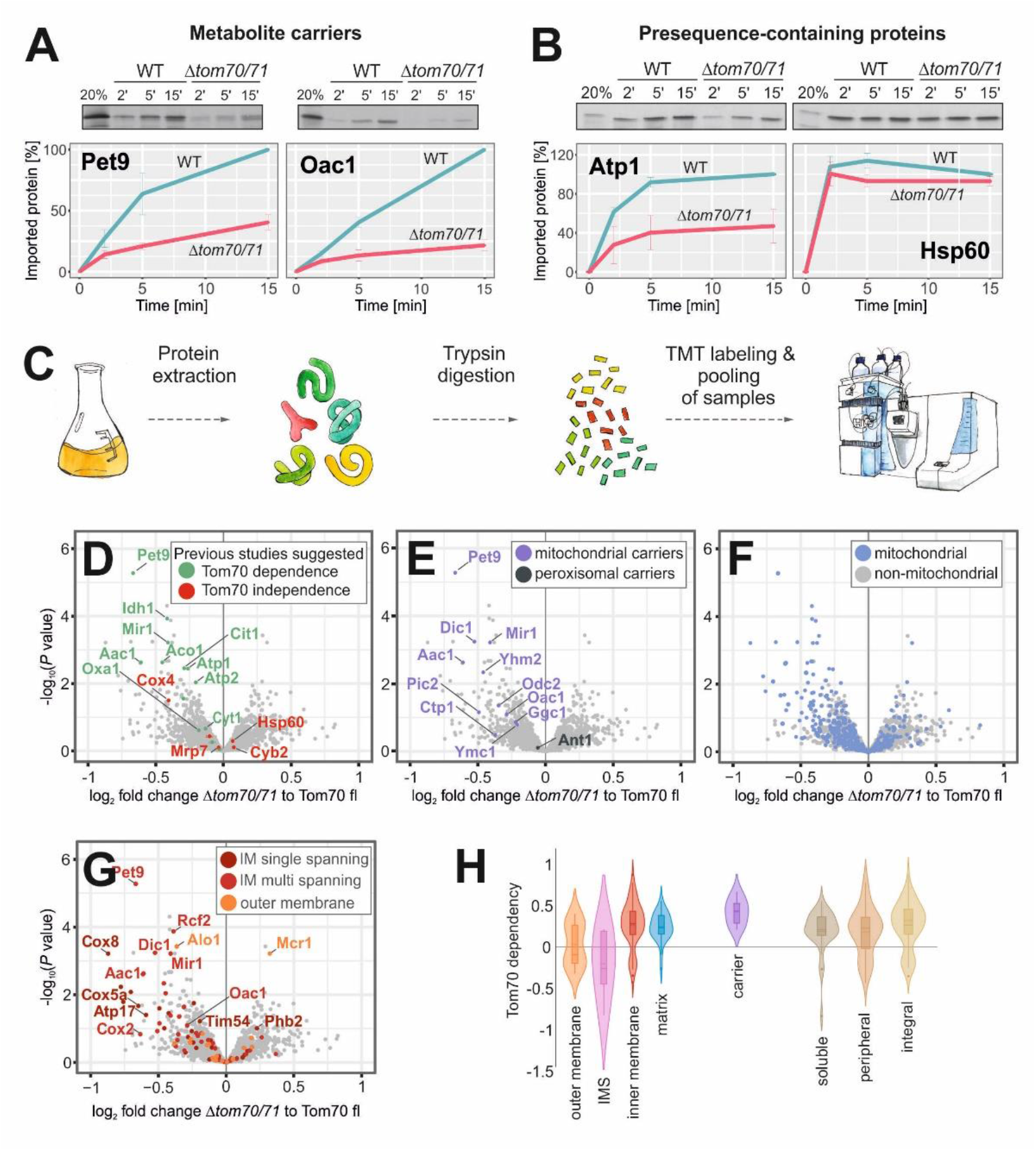
Identification of Tom70/71 clients. **A, B.** Radiolabeled Atp1, Hsp60, Pet9 and Oac1 were incubated with isolated wild type and *Δtom70/71* mitochondria for the times indicated at 25°C. Non-imported protein was removed by treatment with proteinase K and samples were analyzed by SDS-PAGE and autoradiography. Graphs show mean values from three independent experiments. **C.** The proteomes of cell extracts of different mutants were compared using quantitative proteomics and multiplexing. **D-G.** The proteomes of *Δtom70/71* cells carrying either empty or Tom70-expressing plasmids (three replicates each) were measured by mass spectrometry. Shown are the mean values of the ratios obtained from *Δtom70/71* (30°C) to Tom70-expressing cells (30°C) plotted against their statistical significances (*P* values). The points in the upper left corner show the highest Tom70-dependence. Different groups of proteins are indicated in the same data set. IM, inner membrane. **H.** The relative depletion of proteins in the Δ*tom70/71* to Tom70 comparison (log2 fold changes) were taken as proxy for the Tom70-dependence of proteins. Shown are the distributions of these Tom70-dependence values for different groups of mitochondrial proteins (Morgenstern et al., 2017).

This experiment reliably confirmed previous reports about the Tom70-dependent or - independent nature of individual proteins (Fig. 1D). The majority of mitochondrial carrier proteins, but not the peroxisomal carrier Ant1 were reduced in the *Δtom70/71* mutant, albeit to different degrees (Fig. 1E). In particular, Pet9 was confirmed as a Tom70/71-dependent mitochondrial protein, in line with the results from *in vitro* import assays. Interestingly, however, many other mitochondrial proteins were diminished in *Δtom70/71* samples that were never reported to be dependent on Tom70 before (Fig. 1F). This observation points towards a much more general function of Tom70/71 in mitochondrial biogenesis. In particular membrane proteins were found to be affected (Fig. 1G). Many mitochondrial proteins belonging to all types of sub-mitochondrial compartments were consistently, but not strongly reduced in the Δ*tom70/71* samples and none was depleted by more than 40%, explaining the efficient growth of the mutant even on respiratory media.

Taking protein levels in these strains as a proxy for their Tom70-dependence, we found that many inner membrane proteins, including carriers, as well as many proteins of the matrix utilize this outer membrane receptor (Fig. 1H). Proteins of the outer membrane on the other hand showed a heterogeneous Tom70-dependence and most IMS proteins did not show alterations in abundance in Tom70/71 mutants, in consistence with previous studies (Araiso et al., 2019; Gornicka et al., 2014; Lutz et al., 2003). Many soluble proteins were affected in the Tom70 mutants, arguing against a role of Tom70 as a specific receptor for membrane proteins.

### Mitochondrial proteins strongly differ in their dependence on Tom70

As a second, independent strategy to measure the Tom70-dependence of mitochondrial proteins under *in vivo* conditions, we selected a set of 113 MTS-independent mitochondrial proteins N-terminally fused with superfolder green fluorescent protein (sfGFP) and visualized them on the background of Tom70/71 mutants. Specifically, we picked the strains from the N-terminal SWAp-Tag (N-SWAT) library (Weill et al., 2018) in which each of these proteins is tagged with N-terminal sfGFP under the control of a *NOP1* promoter (Fig. 2A, see methods for details). We used fluorescence intensity as a proxy for protein abundance and in addition to determine whether the proteins localize correctly to mitochondria. Using automated approaches, we introduced the tagged strains into genetic backgrounds that lack either Tom70 (Δ*tom70*), its paralog Tom71 (Δ*tom71*), or both (Δ*tom70/71*). High content microscopy screening of these mutants showed three patterns (Fig. 2B-D). The mitochondrial accumulation of some proteins, such as the ATP/ADP carrier Pet9, was considerably reduced in the absence of Tom70 and also affected to a certain degree in *Δtom71* single mutants (Fig. 2B). Some proteins, such as the phosphate carrier Pic2, were Tom71-independent, however, required Tom70 for efficient accumulation in mitochondria (Fig. 2C). A third group, including the dicarboxylate carrier Odc2, was only mildly affected in all mutants, even in the *Δtom70/71* double deletion (Fig. 2D). These results showed that Tom71 supports the biogenesis of some proteins, but is not essential for any of them. Also the relevance of Tom70 was surprisingly variable and even many carrier proteins efficiently accumulated in mitochondria in *Δtom70/71* mutants (Fig. 2D, E).

**Figure 2.**
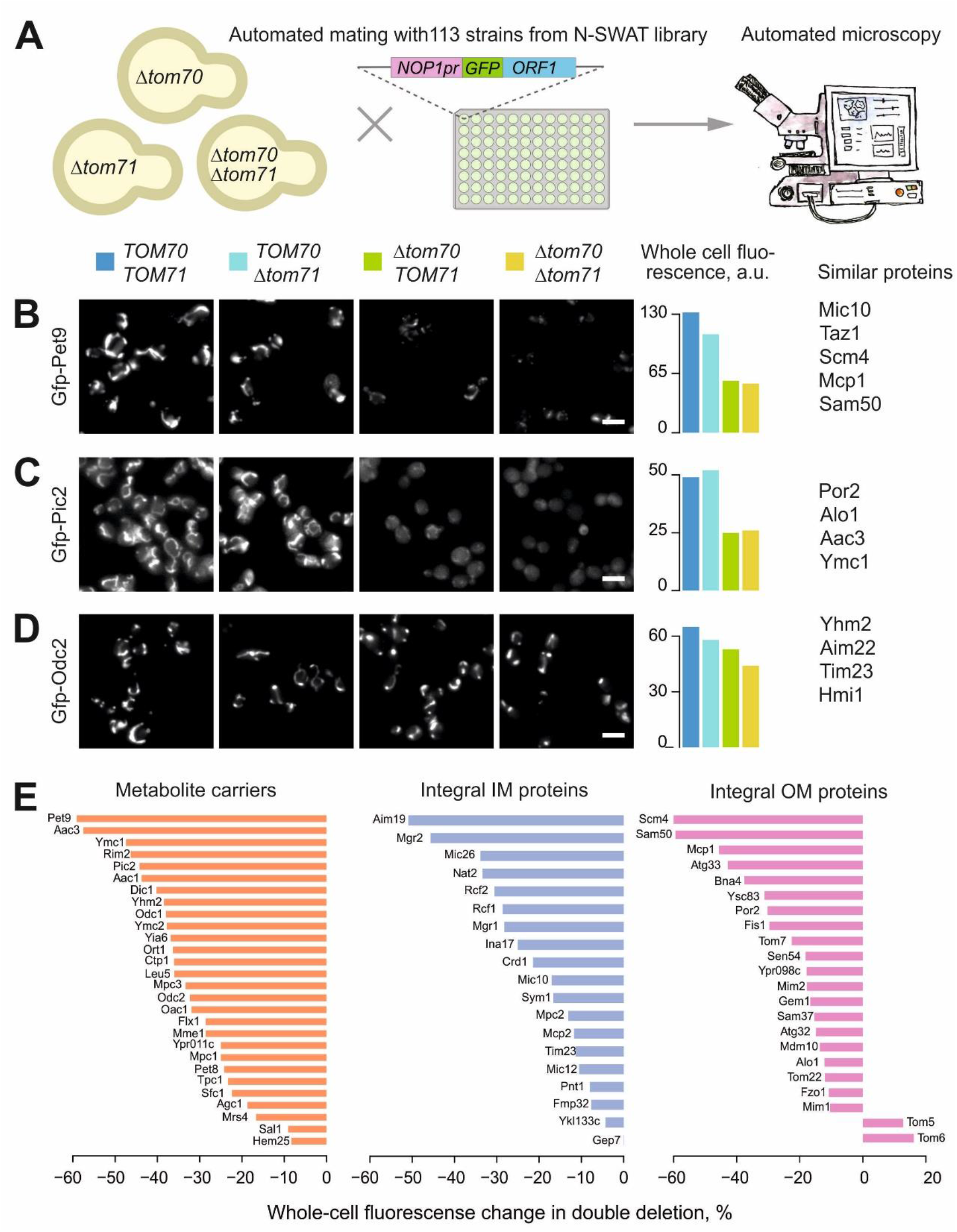
Mitochondrial proteins strongly differ in their Tom70-dependence. **A.** Scheme of the systematic visual screen of GFP-tagged mitochondrial proteins. **B.-D.** The mitochondrial localization of 113 N-terminally GFP tagged mitochondrial proteins (all lacking an MTS) were visualized. Proteins shown in B showed a strongly reduced mitochondrial localization in the absence of Tom70 and moderately reduced levels if Tom71 was deleted. Thus, these proteins depend to some degree on both receptors. Proteins shown in C were unaffected if Tom71 was deleted but still required Tom70. For proteins shown in D, Tom70 and Tom71 were hardly, if at all, relevant. **E.** The whole-cell GFP signal change in Δ*tom70* Δ*tom71* compared to wild type cells measured for different mitochondrial protein classes. Scale bars 10 μm. IM, inner membrane; OM, outer membrane.

By using two orthogonal approaches, we consistently found that, *in vivo*, carriers strongly vary in their Tom70-dependence. The results of the image-based screen with GFP-tagged carriers thereby correlated well with the proteomic data. Carriers are not *per se* more affected in *Δtom70/71* mutants than other mitochondrial membrane proteins. These results challenge the long-standing notion that Tom70/71 is a specialized carrier receptor. Rather, Tom70 and (to a much lesser degree) Tom71 support mitochondrial biogenesis of many proteins. However, the individual dependence of mitochondrial proteins on these receptors varies considerably.

### Tom70/71 supports biogenesis of aggregation-prone mitochondrial proteins

We mined our dataset of Tom70-dependent proteins for features that could determine whether a protein needs the assistance of Tom70 for its biogenesis. As expected, we found Tom70-dependent proteins to be enriched with iMTS-Ls, internal stretches that structurally mimic presequences (Fig. 3A). These were reported before to be efficient binding sites for Tom70 (Backes et al., 2018). In addition, we found significantly higher aggregation propensities for Tom70-dependent proteins (Fig. 3A). The large, hydrophobic carriers are predicted to be particularly aggregation-prone, which might explain why many carriers depend on Tom70 while there is more heterogeneity for other inner membrane or matrix proteins (Fig. 3B). The notion that Tom70 might be required for the correct biogenesis of proteins that are prone to misfold and potentially aggregate is in line with the temperature-sensitive phenotype of the Δ*tom70/71* mutant (Fig. 3C).

**Figure 3.**
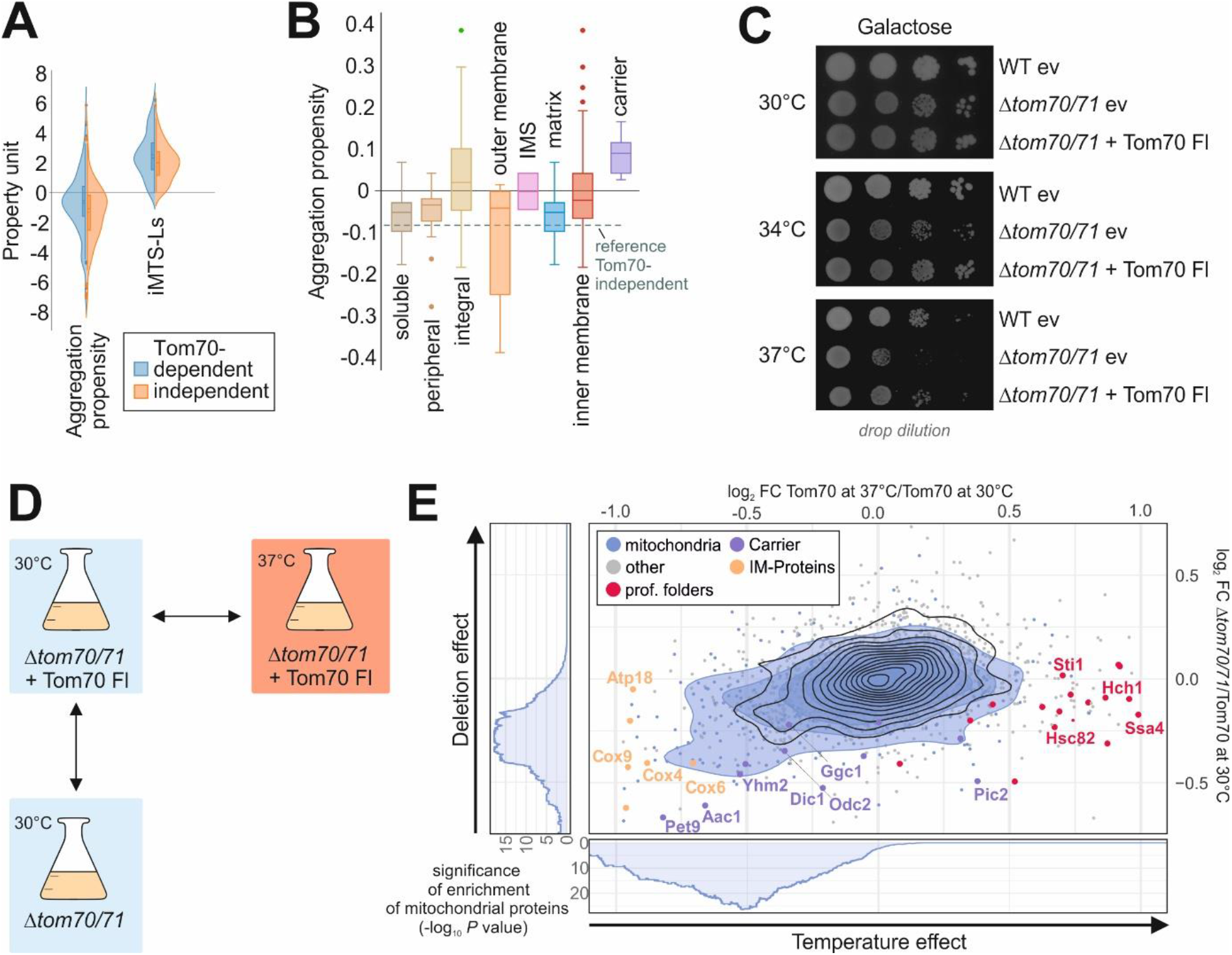
Tom70/71 supports biogenesis of aggregation-prone mitochondrial proteins. **A.** The aggregation propensities (Conchillo-Sole et al., 2007) and the presence of iMTS-L sequences in proteins (Boos et al., 2018) were calculated. Plotted are the distributions of these values for Tom70-dependent (log2 fold change (FC) < −0.2) and independent (log2 FC > 0.2) proteins. **B.** Aggregation propensities were calculated for different groups of mitochondrial proteins. The dotted line shows the mean value of Tom70-independent proteins as reference. **C.** The indicated strains were precultured in galactose-containing medium at 30°C and spotted on galactose medium, following 3 days of incubation at 30, 34 or 37°C. WT, wild type; ev, empty vector. **D, E.** The influence of temperature (log2 FC of Tom70 37°C as compared to Tom70 30°C) and the absence of Tom70 (log2 FC of Δ*tom70/71* as compared to Tom70 at 30°C) were analyzed. Blue circles show the isobaric distribution of mitochondrial proteins, whereas black ones show the distribution of the entire proteome. Enrichment of mitochondrial proteins among proteins with log2 fold change below a certain threshold was calculated and significance of this enrichment was plotted (side panels).

If Tom70 indeed predominantly supports the biogenesis of delicate, aggregation-prone proteins, one would expect that deletion of this stabilizing factor has similar effects on its clients as conditions which promote the general misfolding of proteins. To test this hypothesis, we compared the effect of the absence of Tom70 to that of higher temperature. We grew cells on galactose at 30°C and 37°C and compared the temperature-dependent changes in proteome composition with those caused by deletion of *TOM70/71* (Fig. 3D, E). We observed that many chaperones were found at higher levels in 37°C-grown cells but hardly influenced by the absence or presence of Tom70 (Fig. 3E, marked in red), showing that Δ*tom70/71* cells do not suffer from generally perturbed protein homeostasis. Interestingly, levels of many mitochondrial proteins were reduced by high temperatures as well as by the loss of Tom70, and the effects in these conditions are remarkably similar (Fig. 3E). This is most impressive for Pet9 which was strongly reduced under both conditions, again indicating its aggregation-prone nature.

Hence it appears that it is the tendency to misfold or even aggregate which determines whether a mitochondrial protein requires Tom70 for its biogenesis. This suggests that *in vivo*, the predominant role of Tom70 is not so much that of a receptor that confers directed or specific targeting of its cargo to mitochondria. Rather, it safeguards proteins that are intrinsically prone to acquire non-productive, presumably import-incompetent or even toxic conformations.

### Tom70 can be replaced by a chaperone-tether on the mitochondrial surface

How does Tom70 support the biogenesis of proteins that are prone to misfolding and aggregation? The large size and the multi-domain structure of Tom70/71 indicate that it is not just a simple binder for precursor proteins, as is the case for the much smaller Tom20. Tom70 is tethered to the outer membrane by an N-terminal transmembrane domain and exposes three soluble domains to the cytosol, called Clamp, Core and C-tail domains (Fig. 4A) (Brix et al., 2000; Chan et al., 2006; Young et al., 2003). For simplicity, we refer to them here as C1, C2 and C3. All three domains are formed by tetratricopeptide repeats (TPRs). C2 and C3 were reported to bind internal targeting signals of precursor proteins, such as Pic2 (Brix et al., 2000) (C2) or Adh3 (Chan et al., 2006) and rat alcohol dehydrogenase (Melin et al., 2015) (C3). In contrast, C1 forms a binding groove to recruit the C-terminal EEVD tetrapeptides of Hsp70 and Hsp90 chaperones (Li et al., 2009; Young et al., 2003). We reasoned that it might be this chaperone-binding activity that contributes to the function of Tom70 as stabilizer of aggregation-prone proteins. Expression of a mutant version of Tom70 that only contains the C1 domain proved to be impossible because it was unstable *in vivo* and never accumulated to detectable levels. However, TPR domains are found in many cellular proteins where they serve as specific binding modules for different groups of ligands (Perez-Riba and Itzhaki, 2019).

**Figure 4.**
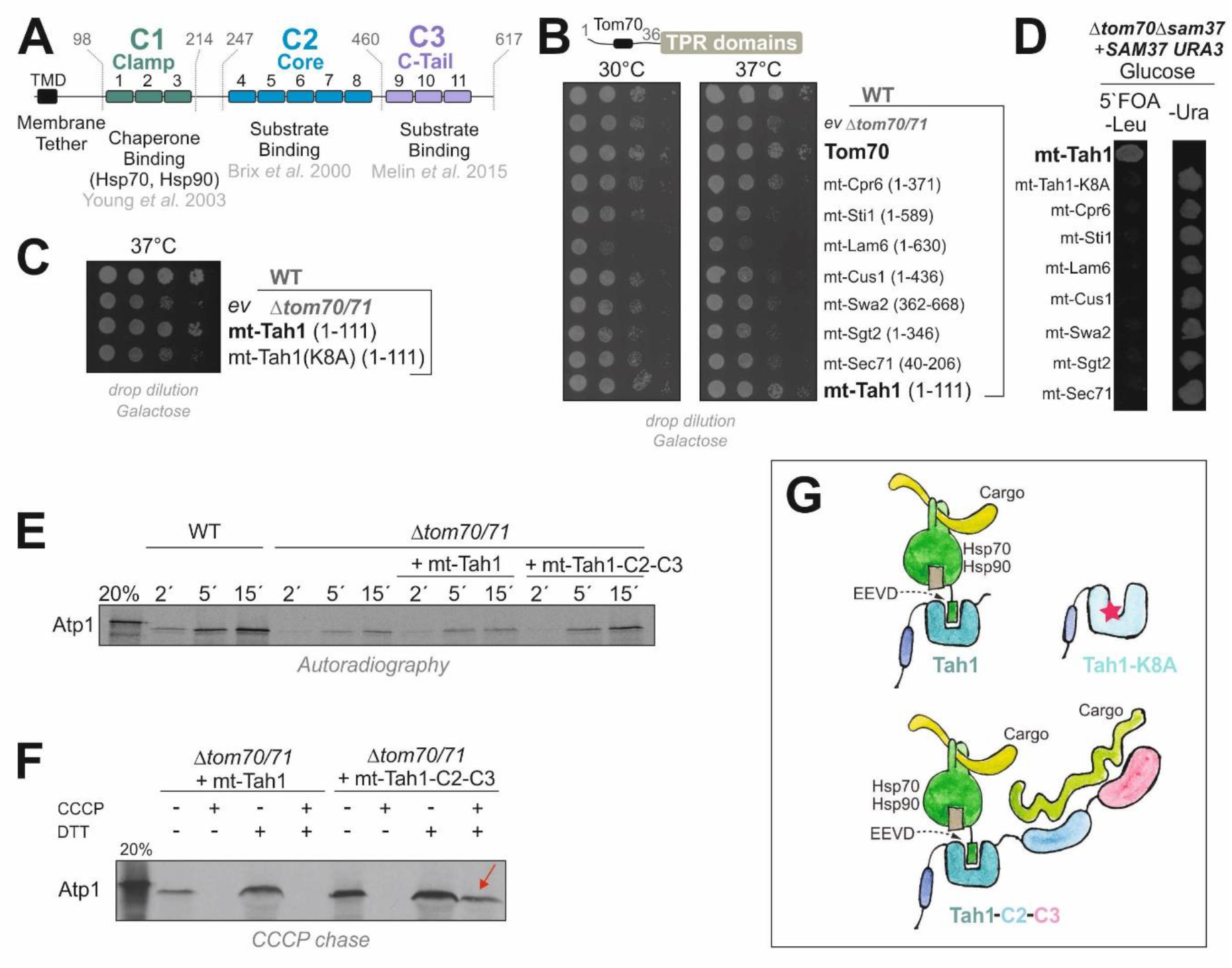
Tom70 can be replaced by a chaperone-tether on the mitochondrial surface. **A.** Schematic representation of the different domains of Tom70 formed by 11 TPR-domains. **B, C.** The indicated sequences of yeast TPR proteins were fused to the membrane anchor of Tom70 and expressed in the *Δtom70/71* mutant. **D.** A *Δsam37 Δtom70* double mutant carrying *SAM37* on a *URA3*-containing plasmid was transformed with plasmids for the expression of the indicated fusion proteins. Upon addition of 5-fluoroorotic acid (5’FOA), only cells which lost the *URA3*-containing *SAM37* plasmid can grow. **E.** Radiolabeled Atp1 was incubated with mitochondria isolated from the indicated mutants. Non-imported Atp1 was removed by adding proteinase K after the times indicated. Mt-Tah1-C2-C3 is a fusion protein in which the C2 and C3 domains of Tom70 were fused to mt-Tah1. **F.** Radiolabeled Atp1 was incubated with mitochondria after the membrane potential was depleted by treatment with carbonyl cyanide *m*-chlorophenyl hydrazone (CCCP). When indicated, CCCP was quenched by dithiothreitol (DTT) to restore the membrane potential. The presence of the C2-C3 domains was essential to keep Atp1 bound to the mitochondria (indicated by the red arrow). **G.** Model of the chaperone binding property of the C1 domain of Tom70/71 and of mt-Tah1. The C2 and C3 domains facilitate direct substrate binding which is particularly relevant under the conditions of the *in vitro* import reaction.

Alignments of the first three TPR domains of Tom70 show sequence similarity to other TPR proteins of yeast which inspired us to test whether any of these other TPR domains can functionally replace Tom70. To this end, we fused six TPR domains to the transmembrane domain of Tom70 and expressed these sequences in the *Δtom70/71* mutant. Tethering the protein Tah1 to the outer membrane (mt-Tah1) suppressed the temperature-sensitive growth of the mutant, whereas the other TPR domains had no or even negative effects (Fig. 4B).

Intriguingly, Tah1, just like Tom70, is an Hsp70/90-binding protein although it functions in a completely different context: Tah1 is a subunit of the cytosolic Rvb1-Rvb2-Tah1-Pih1 (R2TP) assembly complex which facilitates the biogenesis of RNA polymerases and other RNA-binding complexes (Back et al., 2013; Boulon et al., 2010; Jimenez et al., 2012). Its 111 residues form one globular domain comprising two TPR stretches and a capping helix that specifically bind the EEVD-tail of cytosolic Hsp90 (Millson et al., 2008). A point mutation in mt-Tah1 (K8A) which destroys its Hsp90-binding ability (Jimenez et al., 2012) was unable to suppress the phenotype of the *Δtom70/71* mutant (Figs. 4C). mt-Tah1, but not mt-Tah1(K8A) was even able to rescue the synthetic lethal Δ*tom70 Δsam37* double mutant, demonstrating that mt-Tah1 can specifically replace Tom70 and not only indirectly buffers the adverse effects of its absence (Fig. 4D).

In contrast to the *in vivo* situation, in *in vitro* import experiments, mt-Tah1 was not able to replace Tom70 in its ability to facilitate the import of Tom70-dependent substrate proteins such as Atp1 (Fig. 4E). Only when mt-Tah1 was extended with the C2 and C3 domains of Tom70, it was able to support the import of radiolabeled Atp1. This was particularly apparent in ‘CCCP chase experiments’ (Backes et al., 2018; Haucke et al., 1995) in which Atp1 was initially bound to de-energized mitochondria and subsequently chased across the outer membrane upon restoration of the membrane potential (Fig. 4H). In that setup, Tom70 is crucial to ‘hold’ Atp1 on the mitochondrial surface, a function which obviously is not carried out by mt-Tah1. These results confirm that mt-Tah1 rescues the Δ*tom70/71* mutant by replacing it as a chaperone-recruitment factor on the outer membrane (Young et al., 2003), and not by direct binding to precursor proteins (Fig. 4G). In summary, we conclude that the ability of Tom70 to bind substrates directly is largely dispensable under physiological *in vivo* conditions.

### Chaperone-binding by Tom70 is important for different cellular activities

If indeed it is the co-chaperone activity of Tom70 that is essential for cellular and mitochondrial function, we would expect this to be evident from the cellular effects of its absence. To map such global effects, we set out to identify synthetic lethal or sick genetic interactions in Δ*tom70* mutants. We carried out a genome-wide genetic interaction screen (Baryshnikova et al., 2010; Tong and Boone, 2007) by crossing a *Δtom70* query strain with the systematic yeast knock-out library in which all non-essential genes were individually deleted. We measured the colony size of the resulting double mutants as a proxy for their fitness. The genes whose deletions led to considerable fitness reduction or death in the Δ*tom70* but not wild type *TOM70* background were regarded as synthetic sick or lethal genetic interactors of Tom70. We identified many genetic interactors, most of which are components relevant for mitochondrial biogenesis (Fig. 5A), such as Sam37, Tom71, Tom7, Tim8, Tim13, Pam17, Mip1 or Afg3. In addition, we found components relevant for peroxisome biogenesis (Pex17, Pex18), for lipid metabolism (Loa1, Elo1, Elo2, Crd1, Psd1), the ERMES complex (Mdm10, Mdm34, Gem1) and cytosolic proteostasis (Pfd1, Hch1). The fitness of cells lacking any of these components dropped considerably in the absence of Tom70.

**Figure 5.**
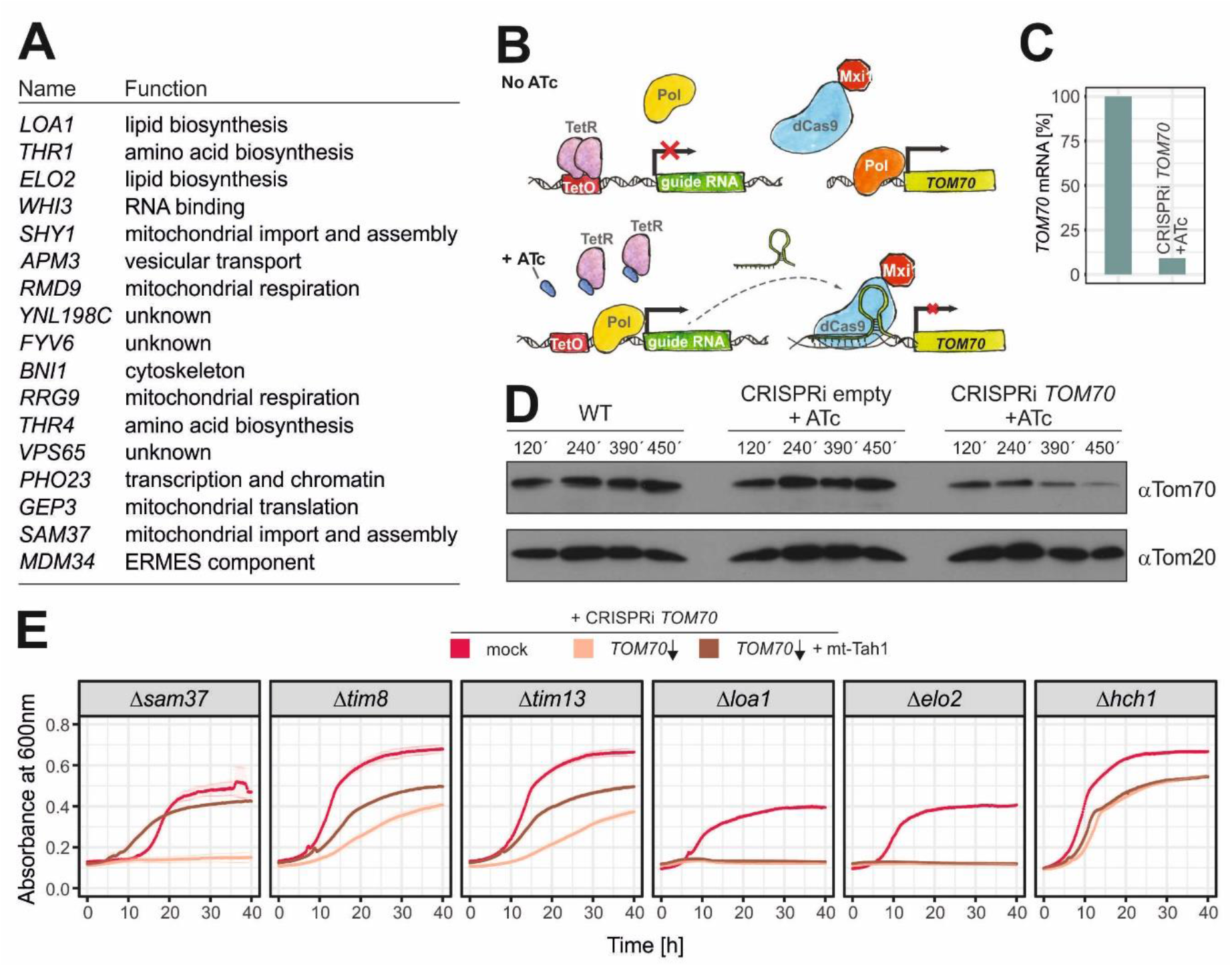
Chaperone-binding by Tom70 is important for different cellular activities. **A.** The *Δtom70* allele was introduced into a systematic yeast deletion library by automated genetic manipulations. Colony sizes were measured and the 100 most affected deletion mutants were analyzed (see Methods for details). **B.** Schematic illustration of the CRISPRi strategy used to knock down *TOM70*. **C**. *TOM70* transcript levels were measured by qPCR 6.5 h after addition of anhydrotetracyclin (ATc). Shown are mean values of three replicates. **D**. Tom70 levels were analyzed by Western blotting of the indicated strains at different time points after addition of ATc. **E**. Growth curves of the following strains: indicated single deletions without addition of ATc (mock); *TOM70* is knocked down through the addition of 960 ng/μl ATc (*TOM70* ↓); *TOM70* is knocked down through the addition of 960 ng/μl ATc, but mt-Tah1 rescues the synthetic growth defect of some mutants (*TOM70* ↓ + mt-Tah1). Shown are mean values and standard deviations from three replicates.

The observation that Tom70 showed genetic interactions with these seemingly different functional groups of proteins could either point to several distinct functions of Tom70 or, alternatively, to the relevance of its chaperone-binding activity for multiple cellular activities. To study these genetic interactions further, despite the lethality of many double mutants, we employed an improved, plasmid-based CRISPRi system to specifically knock down transcription (Fig. 5B). We expressed catalytically inactive Cas9 coupled to a potent transcriptional repressor (dCas9-Mxi1) together with a *TOM70*-specific guide RNA under control of a tetracycline-inducible promoter (Smith et al., 2016). Within a few hours of induction, the levels of *TOM70* mRNA and of Tom70 protein dropped to about 10% (Fig. 5C, D). Combining CRISPRi perturbations with several deletion backgrounds allowed us to verify genetic interaction partners of Tom70, including the genes encoding Tim8, Tim13, Sam37, Loa1, Elo2 and Hch1 (Fig. 5E). Expression of mt-Tah1 in these strains partially rescued the synthetic defects of mitochondrial protein import mutants (Tim8, Tim13, Sam37), but not those in lipid metabolism (Elo2, Loa1). This indicates that the chaperone-recruiting function of Tom70 is of relevance especially in the context of protein biogenesis, while it might be dispensable for other roles.

### Chaperone-binding by Tom70 is crucial for the biogenesis of small inner membrane proteins

Artificial chaperone-recruitment to the outer mitochondrial membrane through mt-Tah1 relieves the temperature-sensitivity of Δ*tom70/71* and its synthetic defects with mitochondrial biogenesis components. To analyze the mechanistic basis of this unexpected observation, we tested how mt-Tah1 influences the proteomic changes that we observed in the Δ*tom70/71* mutant.

Western blots of isolated mitochondria showed that the expression of mt-Tah1, but not that of mt-Tah1(K8A), restores the levels of Tom70-dependent proteins, such as Ugo1, Oxa1 and Pet9 (Fig. 6A, B). Which other proteins rely on the chaperone-binding activity of Tom70? To address this question systematically, we compared the cellular proteomes of Δ*tom70/71* with the same strains expressing Tom70, mt-Tah1 or mt-Tah1(K8A). Indeed, many proteins that were depleted from Δ*tom70/71* cells were rescued by expression of mt-Tah1, but not mt-Tah1(K8A) (Fig. 6C, D). Thereby, mt-Tah1 particularly supported the accumulation of many small proteins of the inner membrane, including many single-spanning subunits of the complexes of the respiratory chain. The levels of these small proteins were reduced in *Δtom70/71* cells but restored if Tom70 or mt-Tah1 were expressed (Figs. 6C-E, protein names labelled in orange).

**Figure 6.**
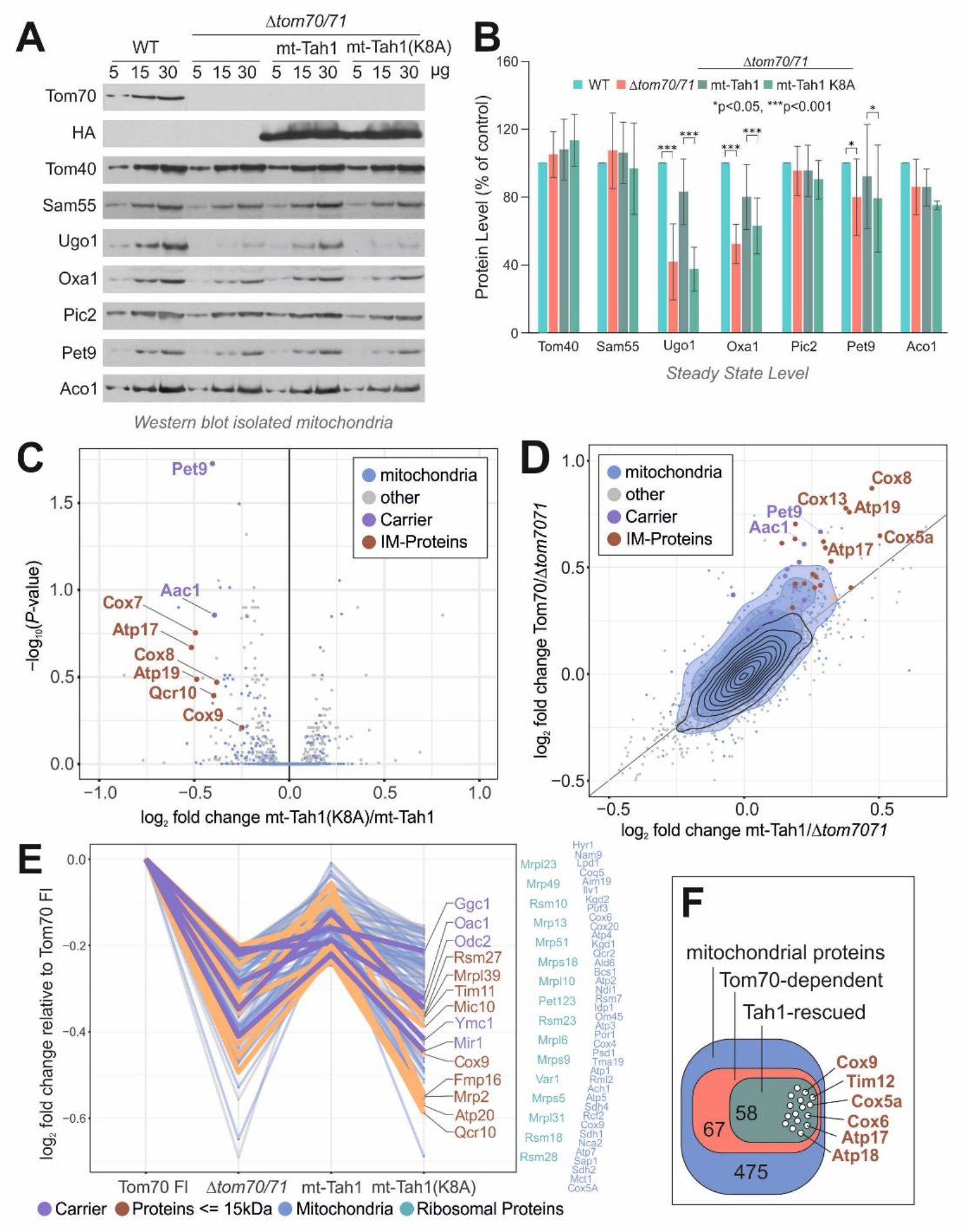
Chaperone-binding by Tom70 is crucial for the biogenesis of small inner membrane proteins. **A, B.** Protein levels in mitochondria isolated from either wild type cells or the indicated *Δtom70/71* mutants were analyzed by western blotting followed by quantification of the relevant bands (n=3). **C.** The volcano plot shows the comparison of the proteomes of *Δtom70/71* cells that express the mt-Tah1(K8A) to those with mt-Tah1. The positions of several small inner membrane proteins (brown) and of carriers (purple), which are considerably stabilized by mt-Tah1 but not by mt-Tah1(K8A) are indicated. **D.** The effects by which Tom70 and mt-Tah1 influence the cellular proteomes are plotted against each other. **E.** Relative log2 fold changes of Tom70-dependent mitochondrial proteins that are rescued by either mt-Tah1 or its variant. **F.** Graphical overview of the number of Tom70-dependent proteins that are rescued by expression of mt-Tah1 near to Tom70 full length levels.

### Chaperone binding by Tom70 prevents mitoprotein-induced toxicity

The inner membrane contains many proteins of less than 18 kDa, most of which are single-spanning non-catalytic subunits of respiratory chain complexes (Morgenstern et al., 2017). These small proteins lack iMTS-L sequences and thus predictable internal binding sites for the C2/C3 domains of Tom70 whereas iMTS-L sequences are ubiquitously found in carriers and other Tom70 clients (Fig. S4A).

The presence of mt-Tah1 restored the levels of many of these proteins (Fig. 7A). Only some of these proteins carry N-terminal presequences, but others do not (Fig. 7B) and their import process was not studied in the past. We therefore chose some of these Tom70-dependent small inner membrane proteins (Cox5a, Tim11, Atp17 and Atp18) as model proteins for further follow-up.

**Figure 7.**
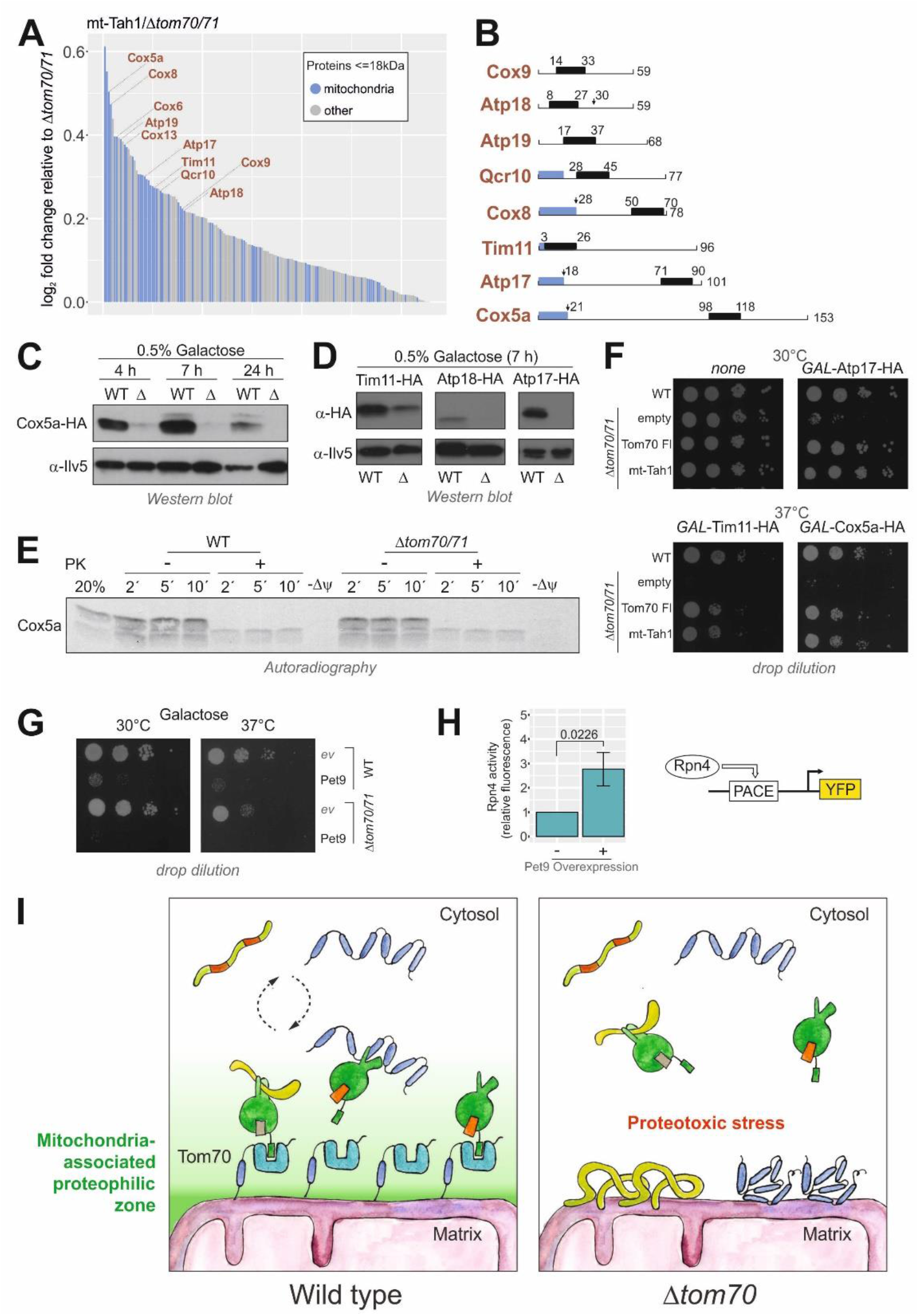
Chaperone binding by Tom70 prevents mitoprotein-induced toxicity. **A.** Proteins that are enriched by the expression of mt-Tah1 in comparison to Δ*tom70/71* (Δ) are shown. Only proteins with masses smaller than 18 kDa and positive enrichment factors were considered. Mitochondrial proteins are indicated in blue. **B**. Schematic representations of small inner membrane proteins for which information about their overall structure and targeting information exists. Blue regions show presequences, black boxes indicate transmembrane domains. **C, D.** Cox5a-HA, Tim11-HA, Atp17-HA and Atp18-HA were expressed under *GAL1* control from multi copy plasmids in wild type and Δ*tom70/71* cells. The times indicate for how long cells were shifted to 0.5% galactose containing medium. **E.** Radiolabeled Cox5a was incubated with isolated mitochondria for the times indicated. The membrane potential (Δψ) was depleted in control samples by addition of CCCP. Mitochondria were reisolated and incubated with or without proteinase K. **F, G.** The indicated strains were transformed with plasmids to express Atp17-HA, Tim11-HA, Cox5a-HA and Pet9-HA under the control of *GAL1* promoter. All cultures were grown on lactate medium to mid-log phase, induced with 0.5% galactose for 4.5 h and dropped onto galactose plates. **H.** Rpn4-driven gene expression was measured using an YFP reporter system (Boos et al., 2019). **I.** Tom70 supports the biogenesis of aggregation-prone mitochondrial membrane proteins by recruiting cytosolic chaperones to the mitochondrial surface, thereby generating a ‘mitochondria-associated proteophilic zone’. See text for further details.

When expressed *in vivo* from a strong *GAL1* promoter, Cox5a accumulated in cells only in the presence of Tom70/71 but not in a deletion mutant (Fig. 7C). The same strong Tom70/71-dependence was observed when Tim11, Atp18 or Atp17 were expressed from *GAL1* promoters (Fig. 7D).

Cox5a was *in vitro* imported into wild type and *Δtom70/71* mitochondria with similar, rather low efficiency (Fig. 7E). This indicates that the striking Tom70-dependence *in vivo* is not recapitulated in the *in vitro* import assay. Aggregation-prone Tom70 substrates could be toxic in the absence of Tom70/71 impeding their intracellular accumulation. Indeed, we observed a high toxicity of overexpressed Cox5a, Atp17, Tim11 and Pet9 (Fig. 7F, G, S4B) and moderate toxicity of Atp18 (Fig. S4C) in the *Δtom70/71* mutant, while wild type cells were less sensitive. The overexpression of Pet9 strongly induced an Rpn4-driven reporter indicative for problems in cytosolic proteostasis (Fig. 7H). An Rpn4-mediated gene induction was observed as a characteristic element of the mitoprotein-induced stress response in yeast (Boos et al., 2019) but is also triggered by other cytosolic proteotoxic stress conditions.

In summary, we conclude that many small inner membrane proteins as well as carrier proteins have the potential to be toxic to cells. Tom70/71 suppresses this toxicity and facilitates the accumulation of these proteins in mitochondria. This property of Tom70/71 depends on its chaperone-binding ability (and hence can be replaced by mt-Tah1). Thus, the primary function of Tom70/71 is that of a cochaperone on the mitochondrial surface that is crucial to protect the cytosolic compartment against proteotoxicity arising from mitochondrial precursors.

## Discussion

Traditionally, the problem of mitochondrial protein biogenesis is seen from the perspective of the organelle. From this direction, Tom70 was identified as ‘ the carrier receptor’, owing to its ability to bind precursor proteins on the mitochondrial surface (via its C2 and C3 domains) and to promote the transfer of these proteins into the protein-conducting channel of the TOM complex. This function of Tom70 is well reflected in the results obtained from *in vitro* import experiments, in which Tom70 clearly supports the import of specific mitochondrial proteins, in particular of the ATP/ADP carrier Pet9 but also of other mitochondrial proteins (Backes et al., 2018; Becker et al., 2011; Brix et al., 2000; Papic et al., 2011; Ryan et al., 1999; Söllner et al., 1990; Wiedemann et al., 2001; Yamamoto et al., 2009). In this study, we employed different approaches to elucidate the relevance of Tom70 in the physiological *in vivo* context. These approaches included the microscopic screens of collections of GFP-tagged proteins, quantitative mass spectrometry-based proteomics of whole cell extracts and the dominant-negative growth effects caused by excess precursor proteins. These three approaches clearly confirmed the particular relevance of Tom70/71 for Pet9 biogenesis. Nevertheless, even though the cellular levels of many mitochondrial proteins, and in particular those of some carriers, were clearly reduced in the absence of Tom70/71, this depletion was never severe. Even for Pet9, the levels in *Δtom70/71* cells were about 40% (GFP signal intensity) to 60% (proteomics) of those in wild type cells, and all other carriers were affected even less. Despite these rather moderate effects, *Δtom70/71* cells are unable to grow under respiration conditions at elevated temperatures. The data presented here point at a crucial function of Tom70 to protect the cytosol against proteotoxic stress conditions for which mitochondrial precursors are presumably responsible. The rather abundant, hydrophobic and aggregation-prone Pet9 protein is particularly problematic here. Previous studies already identified Pet9 as well as its human homolog ANT as proteins with high cytotoxic potential (Coyne and Chen, 2018; Hoshino et al., 2019; Liu et al., 2019; Wang and Chen, 2015; Wang et al., 2008). Mutations that even further increased their aggregation propensity are the cause of Autosomal Dominant Progressive External Ophthalmoplegia 2 in humans (Kaukonen et al., 2000) and trigger mitochondrial precursor over-accumulation stress (mPOS) in yeast (Wang and Chen, 2015). The accumulation of mitochondrial precursor proteins in the cytosol was found to arrest cell growth (Boos et al., 2019; Labbadia et al., 2017; Melber and Haynes, 2018; Wrobel et al., 2015), presumably due to their ability to sequester cytosolic chaperones. Obviously, mitochondrial biogenesis is a considerable challenge for eukaryotic cells (Bar-Ziv et al., 2020; Labbadia et al., 2017) and our observation indicate that Tom70 serves as a component to reduce these mitoprotein-induced stress conditions. In the context of the early stages of precursor targeting to mitochondria, Tom70 apparently serves as the interface between the cytosolic chaperone system and the mitochondrial import machinery (Boos et al., 2020; Hansen et al., 2018; Opalinski et al., 2018; Papic et al., 2013).

How does Tom70 exhibit its cytoprotective activity? In this study, we observe that the artificial tethering of the chaperone-binding TPR protein Tah1 to the mitochondrial surface can almost fully replace Tom70/71 in its properties to promote cell growth at increased temperature, to stabilize the mitochondrial proteome and to provide resistance against overexpressed inner membrane proteins. This observation is astonishing as the function of endogenous Tah1 is completely unrelated to mitochondrial biogenesis. The profound suppression of the *Δtom70/71* mutant by mt-Tah1 indicates that the physiologically relevant function of Tom70 is that of a chaperone-binding factor and that *in vivo*, its direct substrate-binding properties and, hence, its receptor functions are of minor relevance. It appears likely that the recruitment of chaperones to the mitochondrial surface establishes a functionally important ‘mitochondria-associated proteophilic zone’ at the organelle-cytosol interphase that facilitates protein biogenesis and suppresses the potential toxicity of precursor proteins (Fig. 7I). The specific conditions of the *in vitro* import assay where precursors and mitochondria are incubated in a relatively large volume of buffer might have overestimated the relevance of the high-affinity binding sites provided by the C2 and C3 domains of Tom70 for the import of hydrophobic mitochondrial precursor proteins such as carrier proteins.

The comparison of the proteomes of the *Δtom70/71* mutant with that of cells that express mt-Tah1 (chaperone-binding) or Tom70 (chaperone-and substrate-binding) showed that many proteins require the chaperone-binding activity of Tom70, especially those that are intrinsically aggregation-prone. A particularly interesting group of proteins are small (6-18 kDa) inner membrane proteins. Overexpression of these proteins in the absence of Tom70 or mt-Tah1 is highly toxic and prevents cell growth. Apparently, chaperones play a crucial role to facilitate the productive translocation of these proteins to mitochondria. *In vitro*, Tom70 was either not required for these proteins (as for Cox5a) or these proteins could not be imported from reticulocyte lysates (not shown), which points towards a requirement of additional factors – potentially chaperones – that are not accurately reflected in the *in vitro* assay. Finally, we observed that the levels of some Tom70-dependent proteins were not rescued by mt-Tah1. These were particularly inner membrane proteins with bipartite targeting signals such as Sco1 or Dld1. Potentially, these proteins require the direct binding to Tom70 and hence mt-Tah1 does not support their biogenesis. Further studies will be required to study the mechanistic reactions of the Tom70 modules and of cytosolic chaperones in the translocation reactions of these protein classes in more detail.

The cytosol contains many TPR proteins which are structurally similar to Tom70 and Tah1 (Perez-Riba and Itzhaki, 2019; Zeytuni and Zarivach, 2012). Examples are co-chaperones such as Sti1 which, like Tah1 or the C1 domain of Tom70, binds cytosolic chaperones (Hoseini et al., 2016; Schmid et al., 2012). However, TPR proteins apparently are generally employed for the translocation of proteins across cellular membranes. Examples include Sec71 and Sec72 for secretory proteins, Sgt2 for tail-anchored proteins, Pex5 for peroxisomal proteins or Toc64 for plastid proteins (Chartron et al., 2011; Graham et al., 2019; Harano et al., 2001; Schwenkert et al., 2018; Tripathi et al., 2017). It appears likely that early sorting intermediates in general pose a considerable threat for cytosolic proteostasis, that is countered by multiple TPR proteins on organellar membranes. It will be exciting in the future to study the specific roles of this group of proteins more comparatively, not only in respect of their relevance for protein targeting to their respective home organelle, but for cellular proteostasis and fitness in general.

## Acknowledgements

We thank Vera Nehr and Sabine Knaus for technical assistance. We thank Mikhail Savitski, Per Haberkant and Frank Stein for help with setting up the mass spectrometry workflow. We thank Carina Groh and Carsten Balzer for the help with data analysis, and Amir Fadel for help with the microscopy screens. Additionally, we thank Anna Mariam Schlagowski for providing the *GAL1-Pet9* overexpression plasmid, Janina Laborenz for the Tom70-TMD plasmid and Klaus Pfanner and Ophry Pines for antibodies. This study was supported by funding from the Deutsche Forschungsgemeinschaft (DIP MitoBalance to J.M.H, D.R., and M.S. and HE2803/10-1 to J.M.H.), the Joachim Herz Stiftung (to F.B.) and the Forschungsinitiative Rheinland-Pfalz BioComp (to J.M.H. and F.B.). Y.B is supported by an EMBO postdoctoral fellowship. M.S. is an incumbent of the Dr. Gilbert Omenn and Martha Darling Professorial Chair in Molecular Genetics.

## Author Contributions

S.B. and Y.S.B. designed, cloned and verified the constructs and strains; J.M.H. conceived the project; S.B., L.K., F.B., M.R. and Z.S. carried out the mass spectrometry-based proteomics; S.B., F.B., M.R. and T.M. performed the bioinformatical analysis of the mass spectrometry data; S.B., S.L. and J.Z. carried out the biochemical experiments to test the relevance of Tom70 for mitochondrial protein biogenesis; Y.S.B., C.B., and M.S. designed and performed genetic screens to identify Tom70-dependent proteins; C.J., J.D.S. and L.M.S. developed the CRISPR interference strategy and provided the tools which were used by S.B. and S.L.; S.B., D.R., M.S., F.B. and J.M.H. analyzed the data; J.M.H. and F.B. wrote the manuscript with the help and input of all authors.

## METHODS

### KEY RESOURCES TABLE

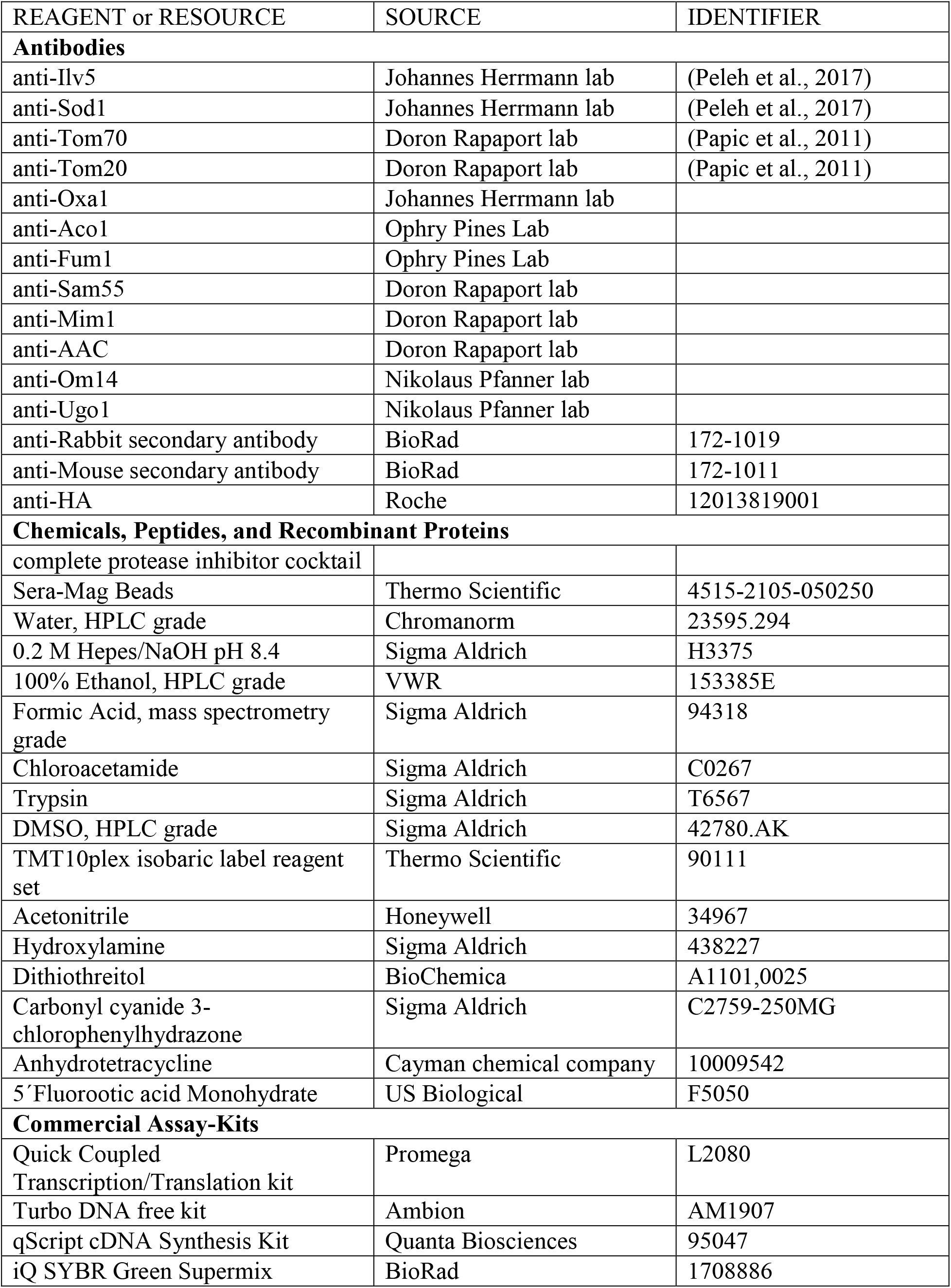
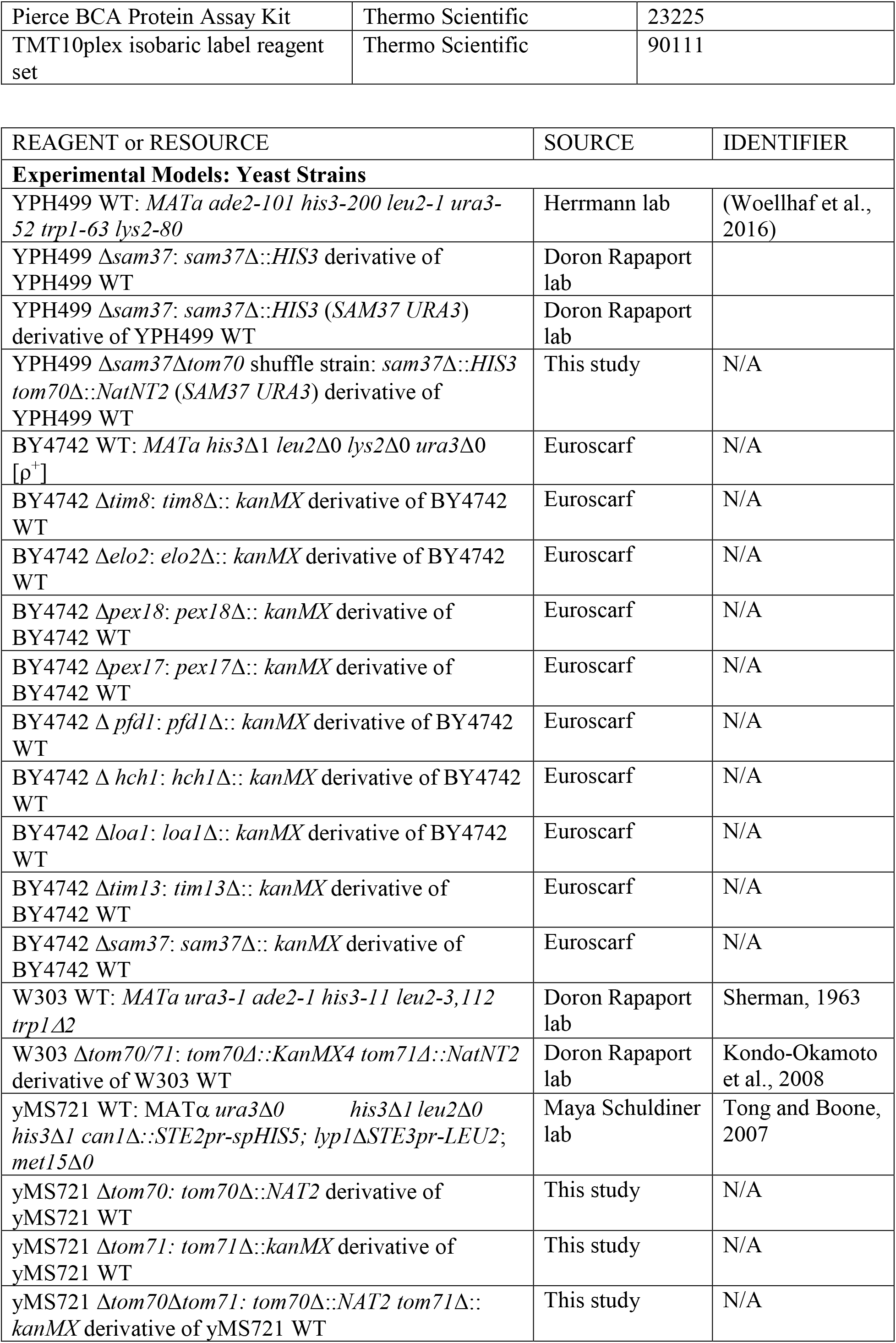
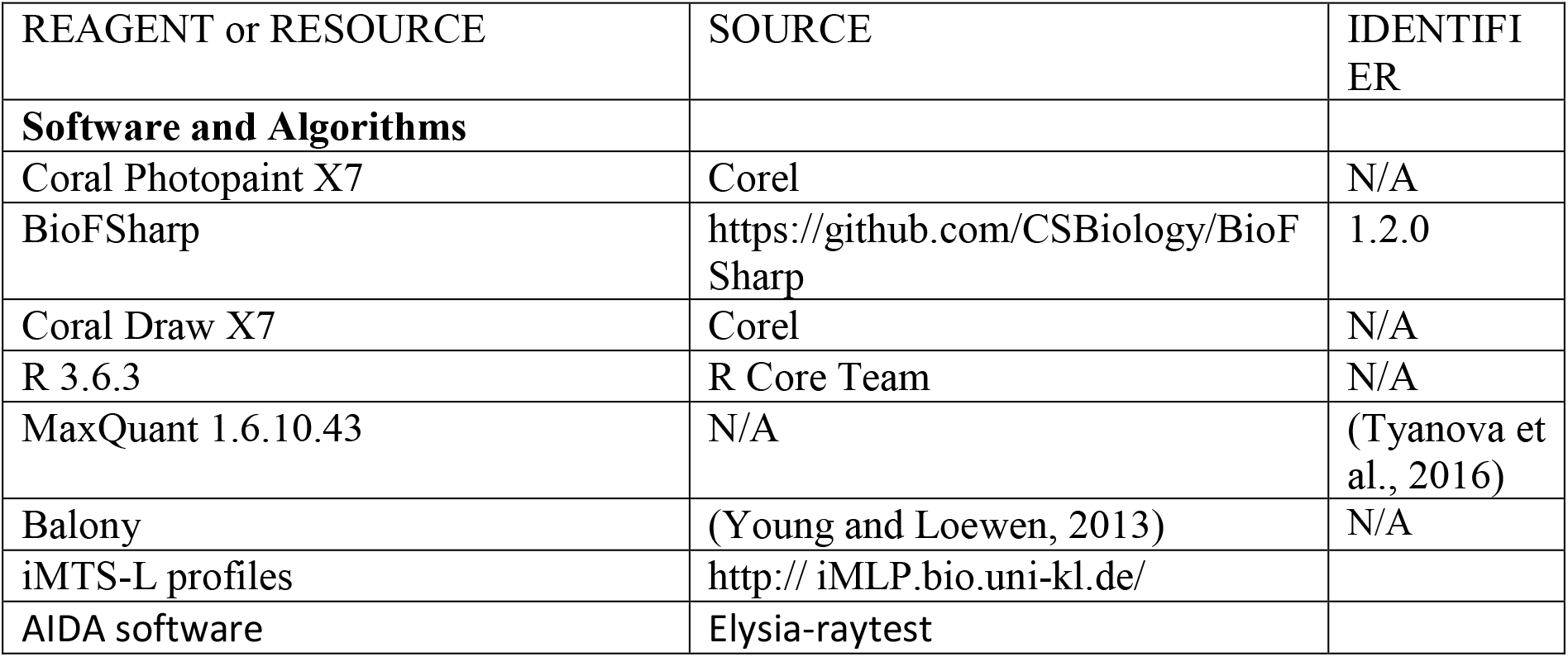

### CONTACT FOR REAGENT AND RESOURCE SHARING

Further information and requests for resources and reagents should be directed to and will be fulfilled by the Lead Contact, Johannes M. Herrmann.

### EXPERIMENTAL MODEL AND SUBJECT DETAILS

#### Yeast strains, plasmids and growth conditions

The yeast strains used in this study are either based on BY4742, W303 or YPH499 background. All strains used in this study are described in detail in the Table above.

All strains were either grown on YP (1% yeast extract and 2% peptone) medium containing 2% glucose or galactose (Altmann et al., 2007) or on minimal synthetic medium (S) containing 0.67% (w/v) yeast nitrogen base and 2% glucose, galactose or lactate as carbon source. To express proteins from the *GAL* promoter, cells were shifted to 0.5% galactose containing medium for 4.5 h.

The shuffle strain for *SAM37* was obtained by replacement of the *SAM37* genomic open reading frame with a *HIS3* cassette in a YPH499 WT. Afterwards, a pRS426-TPI plasmid expressing *SAM37* was transformed, following the subsequent replacement of the *TOM70* genomic reading frame with a *NAT2* cassette.

In order to anchor the TPR-containing proteins to the mitochondrial outer membrane, a pYX142-TPI vector containing the N-terminal Tom70-anchor (residues 1-98) was used to insert the various constructs (for residues see figure 4B). The Tah1 mutation in the MEEVD binding site was achieved by PCR-based site-directed mutagenesis (Quick-Change method, Stratagene) using suitable primer sequences with the desired mutation. For construction of the mt-Tah1-C2-C3 variant, the sequences corresponding to the protein sequence of C2 and C3 domains of Tom70 (residues 247-460 and 461-617) were cloned into the mt-Tah1-containing pYX142 vector.

Yeast transformation was carried out by the lithium acetate method (Gietz et al., 1992). Empty vectors were also transformed in parallel to serve as negative controls.

### METHOD DETAILS

#### CRISPRi system construction

We employ an improved version of a previously generated single plasmid CRISPRi system (Smith et al., 2016) by making it compatible with Type IIS/Golden Gate Assembly, by employing an improved structural gRNA that reduces premature Polymerase III termination (Chen et al., 2013) and by addition of a KanMX resistance cassette. Briefly, we first generated the pKR297 plasmid, containing the RPR1(TetO) promoter, a BspQI-flanking gRNA cassette with an AscI site to remove uncut plasmid, the structural gRNA part and TetR. The pTef1-dCas9-Mxi1-tCyc1 fragment (from pRS416-dCas9-Mxi1: https://www.addgene.org/73796/) was then introduced to yield pKR359 (https://benchling.com/s/seq-gndJVnw6U1oisO0sL65k), and KanMX was inserted to yield pKR366 (https://benchling.com/s/seq-Ymw9j7Wn3MM7g8N7Ny9K).

To assemble the *TOM70* gRNA, two oligonucleotides were annealed in CutSmart buffer (NEB) to form a double-stranded sticky end fragment. pKR366 was digested with BspQI (NEB), gel-purified, and the fragment inserted matching with the BspQI sites using T4 ligase (NEB), according to the supplier’s instructions.

#### Growth assays and viability tests

For spot analysis, the respective yeast strains were grown in liquid rich or synthetic media. Total yeast cells equivalent to 0.5/0.2 OD_600_ were harvested at exponential phase. The cells were washed in sterile water and subjected to ten-fold serial dilutions. Each dilution was spotted on rich or synthetic media followed by incubation at 30°C or 37°C. Pictures were taken after different days of the growth.

Growth curves were performed in a 96 well plate, using the automated ELx808™ Absorbance Microplate Reader (BioTek®). The growth curves started at 0.1 OD_600_ and the OD_600_ was measured every 10 min for 72 h at 30°C. The mean of technical triplicates was calculated and plotted in R. For CRISPRi-mediated repression of *TOM70*, strains were incubated with 960 ng/ml Anhydrotetracycline for 6 h prior to growth curve analysis.

#### High-throughput screening of the GFP collection

To analyze the effect of *TOM70* and *TOM71* deletions on mitochondrial protein import we compiled a mini-library of 113 MTS-independent strains with mitochondrial GFP signal from the N-SWAT-library with NOP1 promotor and N-terminal sfGFP tag, but without a generic MTS inserted before the sfGFP (Weill et al., 2018; Yofe et al., 2016). This mini-library included many members of the metabolite carrier family, other inner membrane proteins, and outer membrane proteins. We didn’t include any MTS-dependent strains in the mini-library since in the N-SWAT library they all carry a very strong generic MTS inserted before sfGFP and thus considerably influencing the import pathway taken by the protein. We constructed Δ*tom70, Δtom71*, and double mutant Δ*Tom70*Δ*tom71* strains in the synthetic genetic array (SGA) compatible background (Tong and Boone, 2007) using standard yeast transformation techniques (Gietz and Woods, 2006; Janke et al., 2004) (See Key Resources Table). We mated these strains with the mitochondrial mini-library and selected for haploid cells harboring both the GFP tag and the required deletion using automated mating and selection approaches as described before (Cohen and Schuldiner, 2011; Tong and Boone, 2007). All mating and selection procedures were performed using RoToR high-density arrayer (Singer Instruments).

For imaging, the resulting haploid libraries with *TOM70*, *TOM71*, or double deletion were inoculated from agar plates into SD-URA liquid media (6.7 g l^−1^ yeast nitrogen base without amino acids, 20 g l^−1^ glucose, and optimized nutrient supplement without uracil according to (Hanscho et al. 2012)) supplemented with 200 μg ml^−1^ nourseothricin for Δ*tom70*, 500 μg ml^−1^ geneticin for Δ*tom71*, or both antibiotics for the Δ*tom17*Δ*tom71* double mutant in 384-well plates and grown overnight at 30°C with shaking. The donor mini-library was at the same time inoculated into SD-URA and grown in the same conditions. All liquid media operations and automated imaging were performed using a JANUS liquid handler (PelkinElmer) connected to an incubator (LiCONiC) and a microscope (Breker et al., 2013). The overnight cultures were diluted 20 times in SD-URA media without antibiotics. After 4 h of growth at 30°C the yeast cultures were transferred to Concanavalin A-coated (Sigma Aldrich) glass-bottom 384-well plates (Matrical Bioscience) and adhered for 20 minutes. Non-adhering cells were washed away with SD-URA-Riboflavin (same as regular SD-URA except yeast nitrogen base without riboflavin and without amino-acids is used to reduce media autofluorescence) that was also used as an imaging media. The plates were transferred to an automated ScanR miscroscopic system (Olympus) using a robotic swap arm (Hamilton). The cells were imaged in bright field and GFP (excitation filter 490/20 nm, emission filter 535/50 nm) channels with 60x air objective (NA 0.9) and the images were recorded on ORCA-ER charge-coupled device camera (Hammamatsu). The donor mini-library and the libraries crossed with *TOM70* and *TOM71* deletion strains were imaged on the same day.

For each strain the four images from the donor library, the library crossed with Δ*tom70*, Δ*tom71*, and Δ*Tom70*Δ*tom71* were displayed side by side and GFP signal localization and intensity were visually assessed.

For quantitive analysis, the cells’ outlines were determined in bright field channel using a custom MATLAB script and median fluorescence intensity was calculated within these outlines. These values were averaged across all detected cells. The strain with the lowest mean cell intensity was taken as a cell background value and this value was subtracted from all other values to obtain background-corrected fluorescence intensities. The background fluorescence for each micrograph was also calculated using background subtraction procedure to assess illumination stability during the imaging process. The illumination was stable throughout the whole imaging period so we performed no additional corrections of the measured cell fluorescence intensities. Mean fluorescence for each strain of the donor mini-library was directly compared to the mean fluorescence of the same strain crossed to *TOM70* and *TOM71* deletions.

#### *TOM70* genetic interaction screen

To investigate genetic interactions of *TOM70* we crossed the Δ*tom70* strain with the yeast full-genome knock-out collection (Giaever et al., 2002) and performed single and double mutant selection as described before (Tong and Boone, 2007). All mating and selection procedures were performed using RoToR high-density arrayer (Singer Instruments).

Briefly, after mating and sporulation all haploid MATα cells were selected on SD-LEU-LYS-ARG (6.7 g l^−1^ yeast nitrogen base without amino acids, 20 g l^−1^ glucose, complete set of supplements without leucine, lysine, and arginine) supplied with canavanine and thialysine. Then the haploids were plated on SD-LEU-LYS-ARG supplied with canavanine, thialysine, and geneticin (G418) to select all haploids that have the library knock-outs disregarding of the *TOM70* allele. From these plates the strains were simultaneously replicated either on the same media (SD-LEU-LYS-ARG + canavanine + thialysine + G418) to measure the library knock out colony size or on the media additionally supplied with nourseothricin (NAT) to select both for the library knock out and Δ*tom70* allele and to measure the colony size of the double mutant. The plates were grown overnight and photographed the next day. Size of the colonies was determined and normalized for each plate using SGAtools (Wagih et al., 2013). For each library knock-out strain the colony size difference between single (library) mutant selection (SD-LEU-LYS-ARG + canavanine + thialysine + G418) and double mutant (library + Δ*tom70}* selection media (SD-LEU-LYS-ARG + canavanine + thialysine + G418 + NAT) was calculated as a measure of genetic interaction with *TOM70* (Table S3).

#### Overexpression assay

Yeast cells were inoculated in non-inducing medium. At mid-log phase (OD 0.6 – 0.8), cells were shifted to inducing conditions (0.5% galactose). At the indicated timepoints, 2 OD_600_ were harvested by centrifugation (20,000 g, 3 min, RT) and whole cell lysates were prepared. Whole cell lysates were prepared for the indicated timepoints to investigate the degradation behavior.

#### Heat-shock assay

Yeast cells were pre-grown in SD medium at 30°C. At mid-log phase (OD 0.6 – 0.8), 0.1 OD_600_ were exposed to a heat-shock at 50°C for the indicated timepoints. After each timepoint, 0.01 OD_600_ were equally plated on SD plates and incubated for two days at 30°C.

#### Sample preparation and mass spectrometric identification of proteins

For mass spectrometry sample preparation, strains were pregrown in S-medium containing 2% galactose at 30°C and either shifted to 37°C or kept at 30°C for 16 h.

50 OD_600_ of cells were harvested at each time point by centrifugation (17,000 g, 3 min, 2°C), washed with prechilled water, snap-frozen in liquid nitrogen and stored at −80°C. Cells lysates were prepared in lysis buffer (50 mM Tris pH 7.6, 5% (w/v) SDS) using a FastPrep-24 5G homogenizer (MP Biomedicals, Heidelberg, Germany) with 3 cycles of 30 s, speed 8.0 m/s, 120 s breaks, glass beads). Lysates were diluted to 2% (w/v) SDS and protein concentrations were determined using the Pierce BCA Protein Assay (Thermo Scientific, #23225). 20 μg of each lysate were subjected to an in-solution tryptic digest using a modified version of the Single-Pot Solid-Phase-enhanced Sample Preparation (SP3) protocol (Hughes et al., 2014; Hughes et al., 2019). Here, lysates were added to Sera-Mag Beads (Thermo Scientific, #4515-2105-050250, 6515-2105-050250) in 10 μl 15% formic acid and 30 μl of ethanol. Binding of proteins was achieved by shaking for 15 min at room temperature. SDS was removed by four subsequent washes with 200 μl of 70% ethanol. Proteins were digested with 0.4 μg of sequencing grade modified trypsin (Promega, #V5111) in 40 μl Hepes/NaOH, pH 8.4 in the presence of 1.25 mM TCEP and 5 mM chloroacetamide (Sigma-Aldrich, #C0267) overnight at room temperature. Beads were separated, washed with 10 μl of an aqueous solution of 2% DMSO and the combined eluates were dried down. In total three biological replicates were prepared (n=3). Each replicate included samples of all 5 strains at 30°C or 37°C (in total 10 samples per replicate). Peptides were reconstituted in 10 μl of H2O and reacted with 80 μg of TMT10plex (Thermo Scientific, #90111)(Werner et al., 2014) label reagent dissolved in 4 μl of acetonitrile for 1 h at room temperature. Excess TMT reagent was quenched by the addition of 4 μl of an aqueous solution of 5% hydroxylamine (Sigma, 438227). Peptides were mixed to achieve a 1:1 ratio across all TMT-channels. Mixed peptides were desalted on home-made StageTips containing Empore C_18_ disks (Rappsilber et al., 2007). The samples were then analyzed by LC-MS/MS on a Q Exactive HF instrument (Thermo Scientific) as previously described.

Briefly, peptides were separated using an Easy-nLC 1200 system (Thermo Scientific) coupled to a Q Exactive HF mass spectrometer via a Nanospray-Flex ion source. The analytical column (50 cm, 75 μm inner diameter (NewObjective) packed in-house with C18 resin ReproSilPur 120, 1.9 μm diameter Dr. Maisch) was operated at a constant flow rate of 250 nl/min. A 3 h gradient was used to elute peptides (Solvent A: aqueous 0.1% formic acid; Solvent B: 80 % acetonitrile, 0.1% formic acid). Peptides were analyzed in positive ion mode applying with a spray voltage of 2.3 kV and a capillary temperature of 250°C. MS spectra with a mass range of 375–1.400 m/z were acquired in profile mode using a resolution of 120.000 [maximum fill time of 80 ms or a maximum of 3e6 ions (automatic gain control, AGC)]. Fragmentation was triggered for the top 15 peaks with charge 2–8 on the MS scan (data-dependent acquisition) with a 30 s dynamic exclusion window (normalized collision energy was 32). Precursors were isolated with a 0.7 m/z window and MS/MS spectra were acquired in profile mode with a resolution of 60,000 (maximum fill time of 100 ms, AGC target of 2e5 ions, fixed first mass 100 m/z).

#### Analysis of mass spectrometry data

Peptide and protein identification and quantification was done using the MaxQuant software (version 1.6.10.43) (Cox and Mann, 2008; Cox et al., 2011; Tyanova et al., 2016) and a *Saccharomyces cerevisiae* proteome database obtained from Uniprot. 10plex TMT was chosen in Reporter ion MS2 quantification, up to 2 tryptic miss-cleavages were allowed, protein N-terminal acetylation and Met oxidation were specified as variable modifications and Cys carbamidomethylation as fixed modification. The “Requantify” and “Second Peptides” options were deactivated. False discovery rate was set at 1% for peptides, proteins and sites, minimal peptide length was 7 amino acids.

The output files of MaxQuant were processed using the R programming language. Only proteins that were quantified with at least two unique peptides were considered for the analysis. Moreover, only proteins that were identified in at least two out of three MS runs were kept. A total of 2920 proteins passed the quality control filters. Raw signal sums were cleaned for batch effects using limma (Ritchie et al., 2015) and further normalized using variance stabilization normalization (Huber et al., 2002). Proteins were tested for differential expression using the limma package for the indicated comparison of strains.

A reference list of yeast mitochondrial proteins was obtained from (Morgenstern et al., 2017). Gene set enrichment analysis was performed using Fisher’s exact test. A Benjamini-Hochberg procedure was used to account for multiple testing, where this was performed (Benjamini and Hochberg, 1995).

#### Data Availability

The mass spectrometry proteomics data have been deposited to the ProteomeXchange Consortium via the PRIDE (Perez-Riverol et al., 2019) partner repository with the dataset identifier PXD021173.

Reviewer account details:

**Password:** shCpyhQn

#### Calculation of aggregation propensity

Aggregation propensity is determined from primary protein sequence using the “hot spot” approach according to N. Sánchez de Groot *et al*. A predictive model is based on the individual aggregation propensities of natural amino acids, which have already been experimentally validated in the literature and provide insights into the effect of disease-linked mutations in these polypeptides. Here, we used the average over a sliding window of 5, 7, 9 or 11 residues depending on total sequence length (<=75, <=175, <=300, or >300). The resulting value is assigned to the central residue in the window and then averaged to obtain the aggregationpropensity of the respective protein (Conchillo-Sole et al., 2007). For convenient application the algorithm was implemented using BioFSharp 1.2.0 (https://github.com/CSBiology/BioFSharp).

#### Miscellaneous

The following methods were performed according to already published methods: Import into isolated mitochondria (Backes and Herrmann, 2017), CCCP chase experiment (Backes and Herrmann, 2017), iMTS-L profile generation (Backes and Herrmann, 2017), isolation of mitochondria (Saladi et al., 2020), whole cell lysates (Saladi et al., 2020), RNA-isolation (Boos et al., 2019), quantitative real-time PCR assays (Boos et al., 2019), PACE-YFP reporter assay (Boos et al., 2019).

